# Synthetic dysmobility screen unveils an integrated STK40-YAP-MAPK system driving cell migration

**DOI:** 10.1101/2021.02.15.431287

**Authors:** Ling-Yea Yu, Ting-Jen Tseng, Hsuan-Chao Lin, Ting-Xuan Lu, Chia-Jung Tsai, Yu-Chiao Lin, Chi-Lin Hsu, Feng-Chiao Tsai

## Abstract

Integrating signals is essential for cell survival, leading to the concept of synthetic lethality. However, how signaling is integrated to control cell migration remains unclear. By conducting a “two-hit” screen, we revealed the synergistic reduction of cell migration when serine-threonine kinase 40 (STK40) and mitogen-activated protein kinase (MAPK) were simultaneously suppressed. Single-cell analyses showed that STK40 knockdown reduced cell motility and coordination by strengthening focal adhesion (FA) complexes. Furthermore, STK40 knockdown reduced translocation of yes-associated protein (YAP) into the nucleus, while MAPK inhibition further weakened YAP activities in the nucleus to disturb FA remodeling. Altogether, we unveiled an integrated STK40-YAP-MAPK system regulating cell migration, and introduced “synthetic dysmobility” as a novel strategy to collaboratively control cell migration.

**One Sentence Summary:** Blocking collaborative pathways within the integrated signaling network synergistically disrupts the migration of cells.

## Main Text

Cell is a robust signaling system. Within it, various pathways interact with each other for survival and cellular functions (*1–3*), creating an impeccable integrated system for signaling. Thus, current antimicrobial and anti-cancer treatments utilized combination therapies to simultaneously disrupt multiple pathways as a practice of “synthetic lethality” (*4–7*). This integrated signaling system is critical for cell survival, yet whether or not it also contributes to cell migration and other cellular functions remains elusive. Moreover, there are extensive studies on cytoskeletal dynamics in cell migration (*8–12*). Yet how different pathways are integrated within the cell signaling network to control cytoskeleton for cell migration remains unclear.

### Two-hit screen to elucidate signaling interaction in cell migration

To address these questions, we designed a “synthetic dysmobility” shRNA-drug two-hit screen to study signaling crosstalk during cell migration in human umbilical vein endothelial cells (HUVECs). In this screen, HUVECs were doubly “hit” by one shRNA and one smallmolecule inhibitor and evaluated by the standardized scratch-wound healing assay (*13*)(Fig. 1A). We first used an shRNA “hitting” one of the 119 genes known to affect cell migration in previously reported screens (*13, 14*). The second “hit” was from one of three drugs inhibiting store-operated Ca^2+^ entry (*15*), Rho-associated kinase (ROCK) (*16, 17*) or mitogen-activated protein kinase (MAPK) signaling (*18*). Concentrations of shRNAs and drugs were carefully titrated to minimize alterations on cell viability (fig. S1A) while their effects on wound healing were maintained (Fig. 1, B and C).

**Fig.1.**
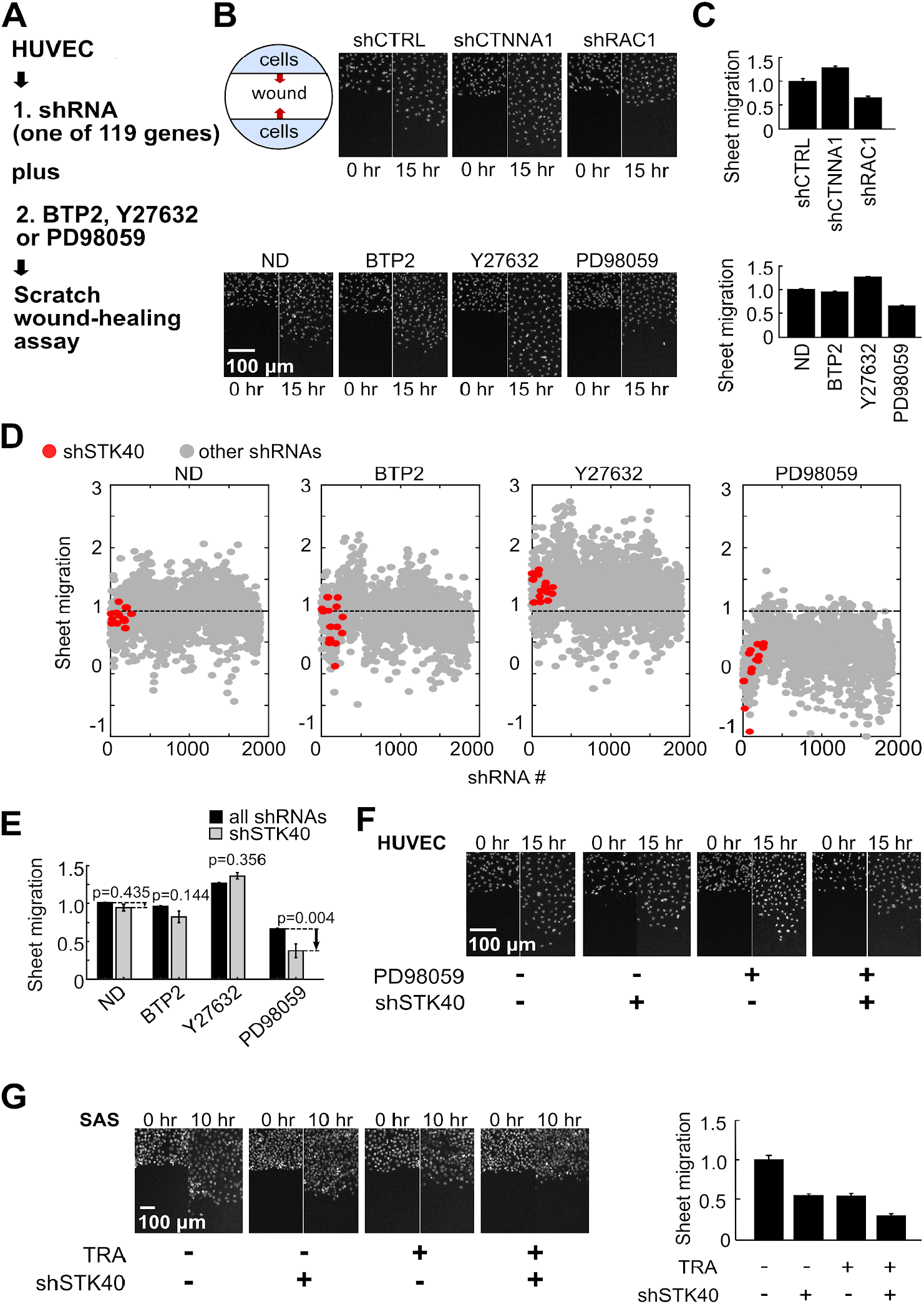
Two-hit screen revealed STK40-MAPK interaction in sheet migration. (A) Scheme for the two-hit sheet migration screen (B) Representative images showing sheet migration of HUVEC cells treated with shRNAs: shCTRL, shCTNNA1 and shRAC1, and drug inhibitors: ND (no drug, *i.e.* DMSO), Ca^2+^ inhibitor BTP2 2 μM, Rho-associated kinase (ROCK) inhibitor Y27632 5 μM, and mitogen-activated protein kinase (MAPK) inhibitor PD98059 10 μM. Hoechst stain marked cell nucleus. (C) Quantification of the sheet migration effects by shRNAs and drugs in (B). (D) Dot plot demonstrates sheet migration effects by individual shRNAs. Red circles resemble shSTK40s while gray circles resemble other individual shRNAs. (E) Black bars resemble average sheet migration of all shRNAs (n=1890), while gray bars resemble average sheet migration of shSTK40 (n=15) upon specific drug treatments. Notice that shSTK40 dramatically decreased sheet migration under PD98059. (F) Representative images of shSTK40 and / or PD98059 -treated sheet migration in our HUVEC screen. (G) SAS cells treated with shSTK40 and another MAPK inhibitor Trametinib (TRA) also revealed synergistic suppression in sheet migration. Left: Representative images. Right: Quantification. Error bars denote mean ± standard error of the mean (s.e.) n=6 biological repeats for each group.

We compared the effects of shRNAs with or without drugs to evaluate their interaction (Data S1) after cell-density correction (fig. S1B) and data normalization (Fig. 1D). When the efficacy of an shRNA was magnified or reduced by a drug, we defined the shRNA-drug interaction to be synergistic (Fig. S1C) or antagonistic (fig. S1D), respectively. Thus, among shRNA-drug pairs, we identified STK40-MAPK interaction (Fig. 1D, and fig. S1, C and D). Previous studies suggested potential involvement of STK40 on MAPK modulation in tissue development (*19*), but its mechanistic linkage remained unclear. We hence began to characterize the role of STK40-MAPK interaction in cell migration. As shown in the screen, shSTK40 together with MAPK suppressor PD98059 synergistically reduced sheet migration in HUVECs (Fig. 1, D to F, and fig. S1C). To verify STK40-MAPK interaction in sheet migration, we further treated both HUVECs (fig. S1E) and head and neck cancer SAS cells (Fig. 1G) with shSTK40 and another MAPK inhibitor Trametinib, and observed synergistic suppression. Therefore, STK40 interacts with MAPK signaling to control sheet migration.

### STK40 changes cell migration by mediating adhesion cytoskeletons

Next, we explored the mechanism underlying STK40-MAPK interaction by focusing on the role of STK40. Since previous screens showed that STK40 knockdown reduced cell-cell coordination (*13, 14*), we analyzed migration behaviors of individual SAS cells by automatic single-cell tracing. Both random migration (Fig. 2A, Movie S1, and Movie S2) and wound healing (fig. S2A, Movie S3, and Movie S4) assays were used because they belong to different modes of cell migration. Cell speed is determined by motility and adhesion in random migration, but by polarity in addition to motility for wound healing. Interestingly, we observed dramatically impaired cell speed and cell-cell coordination by STK40 knockdown in random migration (Fig. 2, A and B). However, in wound healing, cell speed was mildly reduced, and we did not observe consistent change in directionality (fig. S2, A and C). These differential effects of shSTK40 implied that shSTK40 regulated cell adhesion rather than cell migration polarity. Therefore, the role of STK40 is to regulate cell adhesion during cell migration.

**Fig.2.**
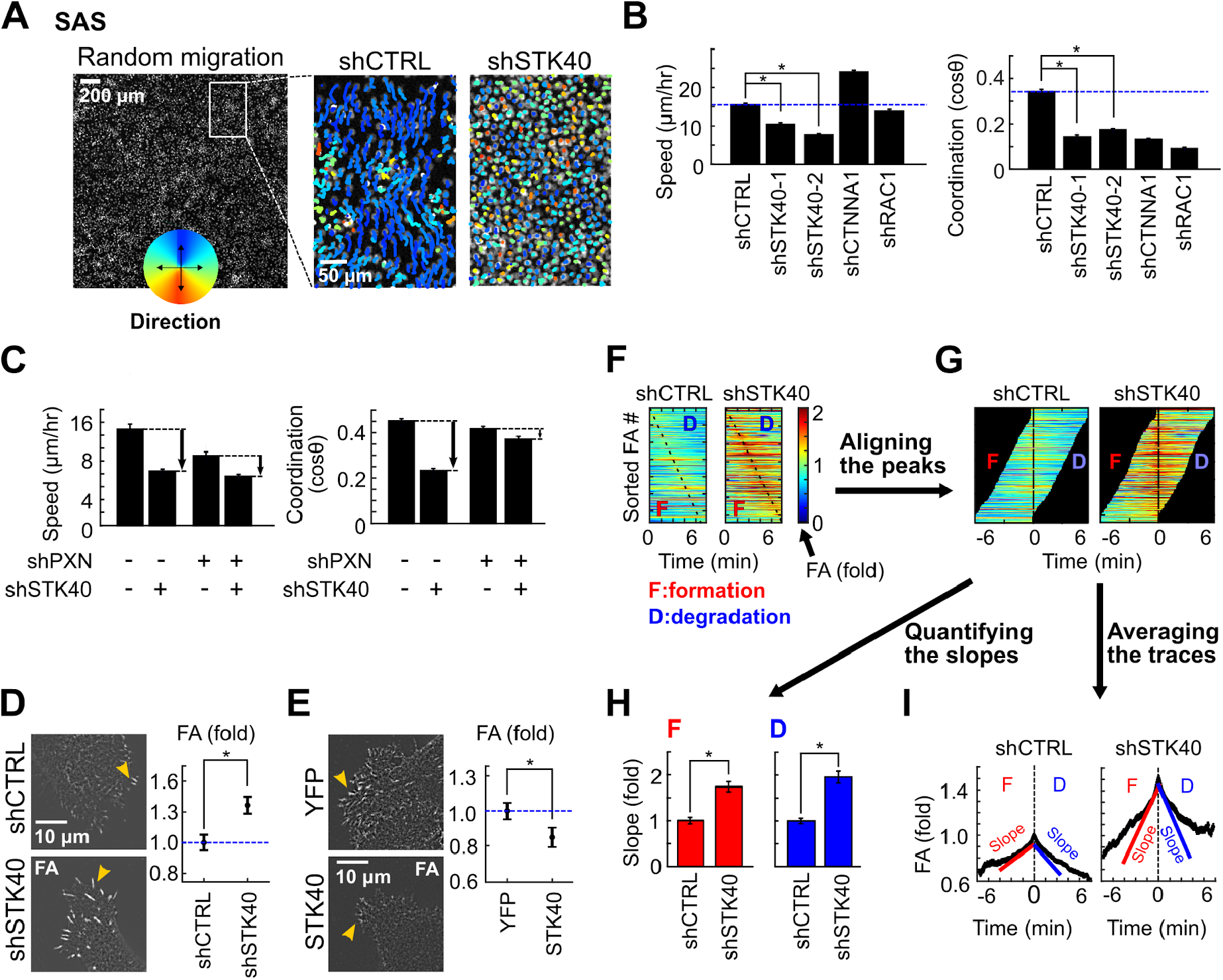
STK40 knockdown reduced cell migration by enhancing focal adhesion (FA) complexes. (A) Random migration experiments in SAS cells. (Left) Cell nuclei were labeled with Hoechst 33342 for tracing of their migration. (Right) Colored lines depicted traces and directions of individual migrating cells treated with shCTRL or shSTK40 within the two-hour period. (B) Well-based quantification showed prominent reduction of cell speed and cell-cell coordination by shSTK40. (C) Double inhibition experiment in random migration showed the reduction of shSTK40 effect by paxillin knockdown (shPXN) in cell speed, and the reversal of the shSTK40 effect by shPXN in coordination. (D and E) Left: Representative images of FA complexes transduced with (D) STK40 shRNAs and (E) control (YFP) and STK40 constructs. Right: Quantification of the integrated FA signals (described in fig. S4E). (F to I) Quantification of FA dynamics using SAS cells expressing GFP-tagged paxillin (PXN). (F) Color maps of FAs sorted by the timing of peak FA signals. (G) Sorted FAs were realigned by the timing of peak signals. (H) Quantification of slopes (derivatives of FA signals over time, depicted in (I)) during FA formation and degradation. (I) Average traces of FA signals based on aligned FA signals in (G). Error bars denote mean ± standard error of the mean (s.e.) \**P* < 0.05.

We further explored the specific adhesion cytoskeleton structure regulated by STK40 with a double-inhibition assay. SAS cells were simultaneously treated with shSTK40 and shRNAs / inhibitors suppressing molecules in the adhesion cytoskeleton structure (Rac1 for Factin (*20*), α-catenin for adherens junction (*21*), paxillin for focal adhesion complexes (*22*, *23*), and myosin light-chain kinase for myosin (*24*))(fig. S2D). We characterized the cell migration behaviors of these treated cells through wound healing (quantified by speed and directionality)(fig. S2, E to H) and random migration (quantified by speed and coordination)(Fig. 2C; and fig. S3, B to E). Interestingly, in both random migration and wound healing, shRAC1 abolished the effect of shSTK40 on cell speed, coordination, and directionality (fig. S2E and fig. S3B) while shSTK40 abolished the effect of ML9 on the same parameters (fig. S2F and fig. S3C). Such epistatic effects implied actin and myosin were both involved in the regulation of STK40-mediated adhesion structure. In contrast, shCTNNA1 showed additive effects to shSTK40 on cell migration parameters (fig. S2G and fig. S3D), indicating the adherens junction was independent from STK40-mediated structure. Strikingly, shPXN not only reduced effects on speed and directionality of shSTK40 (Fig. 2C, and fig. S2H) but also reversed its suppression in coordination (Fig. 2C). This implied paxillin knockdown antagonized the effect of shSTK40. Since paxillin is the major component of the focal adhesion (FA) complex, our results suggested that shSTK40 might disrupt cell migration by strengthening FA complexes.

To answer if shSTK40 enhanced FA complexes, we examined the signals of the anti-paxillin antibody-labeled FA in SAS cells. Indeed, shSTK40 increased both paxillin-labeled FA area and mean paxillin signals overlaying the area (Fig. 2D; and fig. S4), indicating that shSTK40 not only increased the size of FA in two-dimensional manners but also increased the height or strengthened the structure of FA. Moreover, the average number of FA in individual cells was not altered by shSTK40 (fig. S5A), suggesting that STK40 was not involved in the initiation process of FA but was responsible for its turnover. Such STK40 effects on FA were further confirmed by the results that shSTK40 also increased integrated FA signals (total signals from individual FAs) in HepG2 cells (fig. S5B) and HUVECs (fig. S5C), and that overexpression of STK40 reduced integrated FA signals in SAS cells (Fig. 2E). Thus, our results confirmed that shSTK40 disrupted cell migration by strengthening FA complexes.

Next, we explored how shSTK40 altered FA turnover, to see whether shSTK40 strengthened FA by increasing formation or decreasing degradation of FA complexes. We used SAS cells over-expressing EGFP-tagged paxillin in live-cell imaging for the measurement (fig. S5D, and Movie S5) and analyses (fig. S5, E and F; and Fig. 2, F and G) of FA dynamics in SAS cells. STK40 knockdown increased the rates of both FA formation and degradation (fig. S5F, and Fig. 2H) but did not significantly shorten the duration of an average FA turnover cycle (Fig. 2I). Therefore, we inferred that shSTK40 increased FA by facilitating its formation while increased FA degradation probably reflected compensatory responses.

We further studied the mode of action in STK40-FA remodeling using bi-directional approaches. We first dissected the role of STK40 by altering functions and locations of STK40 proteins and examining their effects on FA. STK40 was reported to regulate cell differentiation as a putative nuclear kinase (*19, 25–27*) as well as to maintain protein stability as a cytoplasmic pseudo-kinase (*28, 29*). Thus, we want to know either STK40 location or kinase activity determined FA remodeling. Again, SAS cells overexpressing STK40 protein of full length (FL) had reduced FA (Fig. 3, A and B) and increased cell migration speed (fig. S6A) compared to cells overexpressing the YFP plasmid. In contrast, truncated STK40 with merely its kinase domain (KD) failed to reduce FA. Interestingly, truncated STK40 without its kinase domain (△KD) reduced FA and increased cell speed as the full-length protein did (fig. S6A), indicating that the kinase domain of STK40 was not mandatory for FA remodeling. Moreover, translocating STK40 from the nucleus to the cytosol by tagging it with a nuclear exportation sequence did not significantly alter its reduction effect on FA (Fig. 3, C and D). Together with the facts that shSTK40 did not alter cell size, cell cycle and major markers of epithelial-mesenchymal transition (Fig. S6, B to D), our data suggested that STK40 did not alter FA by regulating cell proliferation or differentiation in the nucleus.

**Fig.3.**
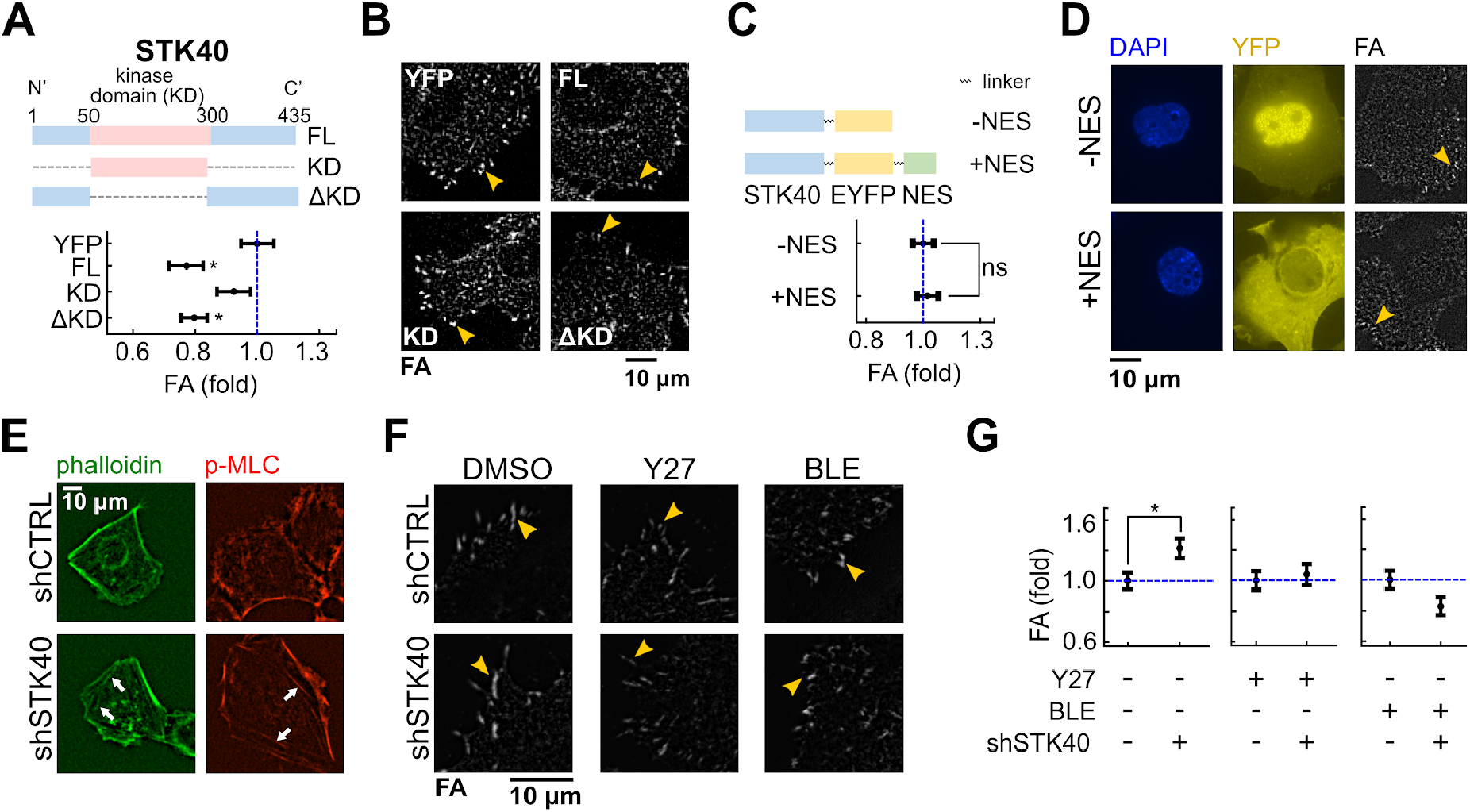
STK40 regulated force-mediated FA formation in kinase-independent manners. (A) Top: Diagram of full-length STK40 (FL), STK40 with kinase domain only (KD), and STK40 with kinase domain-truncated (ΔKD) constructs expressed in SAS cells. Bottom: Quantification of integrated FA intensity in SAS cells over-expressing above vectors. YFP: control vector. (B) Representative images of FAs of (A). (C) Top: Diagram of STK40-EYFP and STK40-EYFP-NES constructs expressed in SAS cells. Bottom: Quantification of integrated FA intensity in SAS cells over-expressing above vectors. (D) Representative images of FA of (C). DAPI channel revealed locations of cell nuclei. YFP channel denotes subcellular distribution of above vectors. Note the distinct cytosolic translocation of STK40 by the NES. (E) shSTK40 increased the number of stress fibers (labeled by phalloidin) and their overlaying phospho-MLC (p-MLC) in SAS cells. (F) Representative images of FA in SAS cells treated with shSTK40 plus DMSO, Y27632 (Y27) 5 μM or blebbistatin (BLE) 5 μM overnight. (G) Quantification of FA in (F). Note the abolishment of shSTK40 effect by both drugs. Error bars denote mean ± s.e. *\*P* < 0.05; ns, not significant.

We then looked into FA and examined if STK40 affected major regulators of FA, including focal adhesion kinase (FAK) signaling (*30–32*) and force-mediated FA strengthening (*33–35*). Since shSTK40 did not change the expression and phosphorylation of FAK (fig. S6E), STK40 probably exerted no direct effect on FAK signaling. Interestingly, when we labeled stress fibers with phalloidin and monitored the contractile activities by overlying phalloidin with the phosphorylated-myosin light chain (p-MLC)(*17*), shSTK40 increased the number of stress fibers and their contractile activities (Fig. 3E). These results indicated that shSTK40 altered FA via force-mediated FA strengthening, which was further supported by the elimination of shSTK40 effects on FA using Rho-associated kinase (ROCK) inhibitor or myosin inhibitor (Fig. 3, F and G; and fig. S6F).

### STK40 collaborates with MAPK on YAP-mediated FA remodeling

Based on the above results, we further asked the question of whether or not MAPK interacted with STK40 for this force-mediated FA strengthening. Indeed, the MEK inhibitor trametinib enhanced FA in SAS cells similarly to shSTK40 (Fig. 4A). Moreover, shSTK40 reduced the effects of trametinib on FA and cell speed, and *vice versa* (Fig. 4A, and fig. S7A). These epistatic effects of trametinib to STK 40 showed that STK40 and MAPK signaling were involved in the same pathway for FA remodeling and cell migration. STK40 was suggested to regulate tissue development by MAPK signalling (*19*, *36*). We also observed that shSTK40 mildly reduced MEK and ERK activities (fig. S7B) and increased the sensitivities in MAPK suppression by trametinib (fig. S7C) while the c-RAF protein levels decreased (fig. S7D). These data supported the role of STK40 in MAPK signaling. However, while shSTK40 only slightly reduced MAPK activities, trametinib completely suppressed it (fig. S7B). Yet both shSTK40 and trametinib induced a marked and comparable increase of FA (Fig. 4A). Such disproportionate results between shSTK40-induced MAPK suppression and FA enhancement suggested the existence of missing links between STK40 and MAPK signaling.

**Fig.4.**
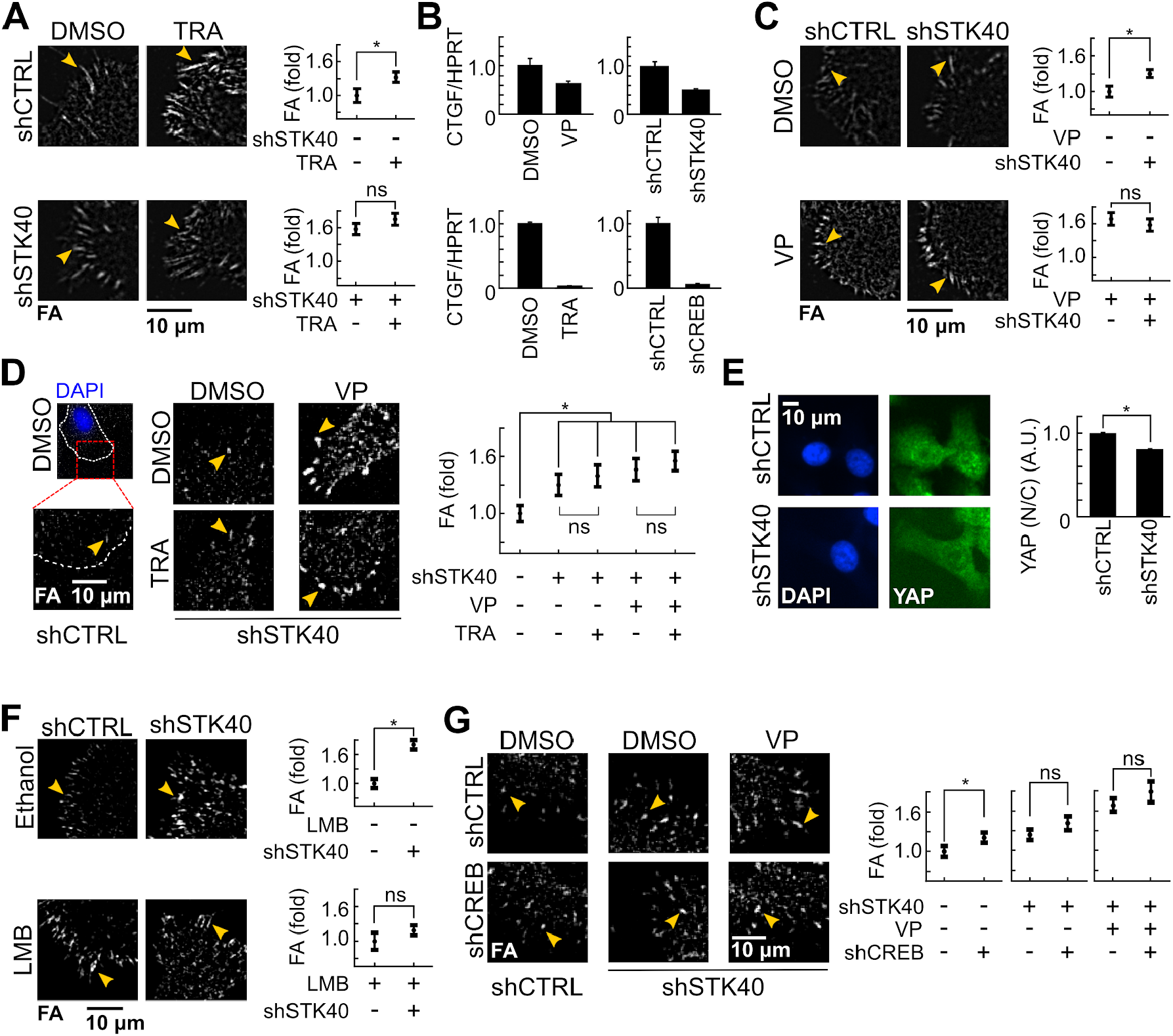
STK40 and MAPK signaling regulated YAP-mediated FA remodeling in collaborative manners. (A) Representative images and quantification of FA in SAS cells without or with shSTK40 treated with trametinib (TRA) 100 nM overnight. Note the vanishment of trametinib effect under shSTK40. (B) YAP activity (labeled by CTGF expression) was decreased in SAS cells treated with shSTK40, shCREB, verteporfin (VP) or trametinib. (C) Representative images and quantification of FA in SAS cells without or with VP treated with shCTRL or shSTK40. (D) Representative images and quantification of FA in SAS cells with shSTK40 and treated with VP and / or trametinib. Note that shSTK40 and VP abolished effects of trametinib. (E) Representative images and quantification of subcellular YAP distribution. Notice the decreased nucleus-to-cytosol (N/C) ratio of YAP by shSTK40. (F) Representative images and quantification of FA in SAS cells without or with leptomycin-B (LMB) treated with shCTRL or shSTK40. Note the vanishment of STK40 effect under LMB. (G) Representative images and quantification of FA in SAS cells with shSTK40 and treated with VP and / or shCREB. Note that shSTK40 and VP abolished effects of shCREB. Error bars denote mean ± s.e. \**P* < 0.05; ns, not significant.

We investigated the missing links. The increase in FA by trametinib was seen 16 hours after treatment (Fig. 4A) but not visible within the first 8 hours (fig. S7E), indicating potential involvement of transcriptional control in MAPK-mediated FA remodeling. Thus, we explored if there existed any transcriptional machinery specifically responsible for FA remodeling. Excitingly, recent reports demonstrated that suppression of the yes-associated protein (YAP) pathway increased FA formation (*37*–*39*) similarly to our results of STK40 knockdown or MAPK inhibition. Such similarities in phenotypes encouraged us to examine if the YAP pathway was involved in STK40-MAPK interaction. Indeed, using the expression of connective-tissue growth factor (CTGF)(*40*) as the readout, we demonstrated that the activity of YAP was suppressed by STK40 knockdown, and also by MAPK suppression (Fig. 4B)(*41*). The YAP inhibitor verteporfin (VP) not only increased FA as reported previously (*39*) but also fully abolished the effects of STK40 knockdown (Fig. 4C) or MAPK inhibition (Fig. 4D). Therefore, we inferred that the YAP pathway took a dominant role on STK40-MAPK interaction for FA remodeling.

Next, we worked on the mechanism of how STK40 and MAPK were involved in YAP-mediated FA remodeling. Western blots suggested that STK40 regulated YAP levels as it remarkably abolished trametinib-mediated compensatory YAP accumulation (fig. S7G). Literature showed that YAP localization was critical for its activity (*42*), as YAP in the nucleus regulated gene expression while YAP in the cytosol was prone to be degraded (*43*). We thus examined if STK40 altered the expression and spatial distribution of YAP at the subcellular level. Intriguingly, immunofluorescence revealed that shSTK40 reduced YAP level both in the nucleus and cytosol (fig. S7F), together with the reduction of its nucleus-to-cytosol (N/C) ratio (Fig. 4E). Such results indicated that shSTK40 might both alter the levels of YAP by reducing its expression or by promoting its degradation, and change the distribution of YAP between the nucleus and cytosol. Previous reports showed that STK40 bound to and activated COP1 (*28*), which functioned as the E3 ubiquitin ligase for protein degradation. Hence, shSTK40 would reduce protein degradation, which was not compatible with its reduction of YAP level in our observation, suggesting that shSTK40 did not directly enhance YAP degradation. In contrast, STK40 knockdown was reported to increase the expression of histone deacetylase 5 (HDAC5)(*25*), which in turn might suppress YAP expression (*44*) and its effect on cytoskeletal remodeling. Moreover, our results about the decreased N/C ratio of YAP by shSTK40 (Fig. 4E) favored the possibility that shSTK40 reduced YAP effect on FA by increasing its cytosolic translocation. This speculation was supported by our finding that leptomycin-B (*45*), which blocked nuclear export to cause YAP accumulation in the nucleus (fig. S7H), abolished shSTK40-induced FA increase (Fig. 4F). Therefore, STK40 probably altered FA remodeling by regulating the expression and subcellular distribution of YAP.

Lastly, we examined how MAPK inhibition regulated YAP effects on FA remodeling. Since both trametinib and shCREB suppressed YAP activity (Fig. 4B) and increased FA (Fig. 4A, and Fig. 4G), we proposed that MAPK signaling augmented YAP effects on FA via its downstream target CREB (*46*). Indeed, similarly to trametinib treatment (Fig. 4D), the FA-enhancing effect by shCREB was also reduced under STK40 knockdown or YAP inhibition (Fig. 4G), confirming the involvement of CREB to augment YAP activities. Though decreasing YAP activity as indicated by CTGF reduction (Fig. 4B), trametinib increased intracellular YAP levels (fig. S7G). Such results indicated that CREB acted downstream of YAP, so MAPK inhibition resulted in compensatory increase of YAP levels. Taken together, we discovered an integrated YAP-centered system regulating cell adhesion and cell migration. Within this system, STK40 promoted nuclear localization of YAP and hence its activity, which was further augmented by MAPK signaling via CREB to control adhesion cytoskeleton. Therefore, simultaneous suppression of STK40 and MAPK collaboratively disrupted FA turnover resulting in “synthetic dysmobility” (fig. S7I).

## Discussion

Previous studies unveiled potential involvement of STK40 on tissue development (*19*, *25–27, 29, 36*), but its molecular mechanism has remained elusive. Our research resolved this issue by confirming the kinase-independent effect of STK40 on force-mediated FA turnover, and by elucidating STK40 regulation on YAP expression and localization. Hence, STK40 likely regulates tissue development via its kinase-independent action on YAP signaling. Our results also identified the missing links between STK40 and MAPK (*19*) by unraveling how STK40 collaborated with MAPK on different levels of YAP control. Via this STK40-YAP-MAPK interaction, we demonstrated how different signaling pathways constituted an integrated system to drive cytoskeletons. First and foremost, our screen disclosed that cell migration could be blocked by multi-targeting strategies mimicking the concept of “synthetic lethality”, which will lead to the development of combination drug therapies to stop cell migration in collaborative manners.

## Acknowledgments

We thank Dr. Tobias Meyer (Cornell University, New York, NY, U.S.A.), Dr. Sean Collins (University of California, Davis, CA, U.S.A.) and Dr. Arnold Hayer (McGill University, Montréal, QC, Canada) for critical discussions about the design of the two-hit screen, Dr. Tzuu-Shuh Jou (National Taiwan University, Taipei, Taiwan), Dr. Keng-Hui Lin (Academia Sinica, Taipei, Taiwan) and RNAi Core (Academia Sinica, Taipei, Taiwan) for technical assistance about the screen, and Dr. Irene Tai-Lin Lee, Ms. Hulda Yu-Chun Chen, Dr. Jessica Yu and Dr. Chia-Ling Ann Yang for critical comments.

## Funding

This work was supported by grants from the Ministry of Science and Technology in Taiwan (MOST 107-2320-B-002-038-MY3), National Taiwan University Hospital (NTUH106-T02, NTUH107-T13, NTUH108-T13 and VN109-14) and the Liver Disease Prevention & Treatment Research Foundation in Taiwan.

## Author contributions

F.-C.T. conceived and designed the two-hit screen, C.-J.T. conducted the screen. F.-C.T. and H.-C.L. analyzed the screen data. L.-Y.Y., T.-J.T., H.-C.L. and T.-X.L. designed and performed most experiments following the screen. Y.-C.L. assisted in the fluorescent microscopy. F.-C.T., L.-Y.Y., T.-J.T., T.-X.L., and C.-L.H performed data analysis and interpretation. F.-C.T. and L.-Y.Y. wrote the manuscript, with assistance from T.-X.L. and C.-L.H. and input from all other coauthors.

## Competing interests

Authors declare no competing interests.

## Data and materials availability

All data is available in the main text or the supplementary materials.

## Supplementary Materials

Materials and Methods

Figures S1-S7

Movies S1-S5

Data S1-S5

## Notes

### Competing Interest Statement

The authors have declared no competing interest.

