## Supplementary Material for "Synthetic dysmobility screen unveils an integrated STK40-YAP-MAPK system driving cell migration"

**This PDF file includes:**

Materials and Methods
Figures S1 to S7
Captions for Movies S1 to S5
Captions for Data S1 to S5

**Other Supplementary Materials for this manuscript include the following:**

Movies S1 to S5
Data S1 to S5

### Materials and Methods

Cell Culture

SAS cells were gifts from Jean San Chia's Lab (National Taiwan University, Taipei, Taiwan). HepG2 and HEK293T cells were purchased from ATCC (Manassas, VA, USA), while HUVEC cells were purchased from Lonza (Lonza, Basel Stücki, Switzerland). Cells were maintained at 37°C and 5% CO2. SAS, HepG2 and HEK293T cells were grown in Dulbecco's modified Eagle's medium (Gibco, Thermo Fisher Scientific, Waltham, MA, USA and HyClone, Logan, UT, USA). HUVEC cells were grown in EGM2 (Lonza). Culture medium was supplemented with 1% of penicillin/streptomycin (Gibco) and 10% fetal bovine serum (HyClone). Cells were passaged using 0.5% trypsin-EDTA (Gibco) and washed with phosphate-buffered saline (PBS) (Corning, New York, USA) within 10 passages.

Generation of knockdown plasmids and overexpression constructs

To knockdown STK40 or other genes (RAC1, CTNNA1, PXN, YAP, and CREB1), we transfected shRNA plasmids bought from National RNAiCore (NRC, Academia Sinica,Taipei, Taiwan) into HEK293T cells to prepare lentiviruses for infection (Data S2). To overexpress STK40, we created 6 different constructs (Data S3).

(1) Control construct pLAS2w.eYFP.Pbsd was created by digesting the vector pLAS2w.Pbsd (Academia Sinica) using restriction enzymes NheI (NEB, MA, USA) and EcoRI (NEB), then ligating cloned YFP using T4 ligase (NEB) to the vector. YFP sequence was cloned from primers *eYFP_NheI* and *eYFP_EcoRI* and pEX-SP-eYFP.STIM1 (gift from Tobias Meyer's lab, Stanford, CA, USA) and was cloned to pLAS2w.Pbsd vector.

(2) pLAS2w.STK40-P2A-EYFP was created by digesting vector pLAS2w.Pbsd using restriction enzymes NheI and XbaI (Takara, Shiga, Kyoto, Japan) and ligating two inserts, STK40 and P2A-EYFP, to the vector. Cloned STK40 section was created by performing nested-polymerase chain reaction (PCR) with two sets of primers: *STK40_NstPCR1/F*, *STK40_NstPCR1/R* and *STK40 cds RE NheI F*, *STK40 cds XbaI R*, using extracted cDNA from HepG2 cells as template. P2A-EYFP was cloned by performing PCR with primers *P2A-EYFP RE XbaI F* and *P2A-EYFP RE SbfI R* using pLAS2w.FGFRIIIb-P2A-EYFP as template.

(3) pLAS2w.STK40-EYFP was generated from pLAS2w.STK40-P2A-EYFP by performing PCR, site-directed mutagenesis and self-ligation. Primers *RE XbaI EYFP-F* and *STK40 cds XbaI R*, template pLAS2w.STK40-P2A-EYFP, and reagents from Quikchange site-directed mutagenesis kit (Agilent, Santa Clara, CA, USA) were utilized for PCR. The enzyme DPN1 from the kit was added to the PCR product to degrade the original template. Restriction enzyme XbaI was further used for digestion of the vector, and self-ligation was performed to create the final construct.

(4) pLAS2w.STK40-EYFP-NES was created by replacing STK40-EYFP from pLAS2w.STK40-EYFP with STK40-EYFP-NES. pLAS2w.STK40-EYFP was digested using restriction enzymes NheI and SbfI. Cloned insert STK40-EYFP-NES was then ligated to the vector. STK40-EYFP-NES was generated by performing PCR with primers *STK40 cds RE NheI F* and *STK40-EYFP-NES primer R pLAS2w.SbfI*, utilizing pLAS2w.STK40-EYFP as template.

(5) pLAS2w.STK40-ΔKD(51-299)-P2A-EYFP (KD: kinase domain) was generated by truncating amino acids 51 to 299 of STK40 from pLAS2w.STK40-P2A-EYFP. PCR was carried out using primers *Kinase domain truncate 300 aa XmaI primer F* and *Kinase domain truncate 50 aa XmaI primer R* and pLAS2w.STK40-P2A-EYFP as template to clone the sequence excluding of the kinase domain. DPN1 (Agilent) was used to degrade template pLAS2w.STK40-P2A-EYFP. Restriction enzyme XmaI (NEB) was used for digestion and self-ligation was then performed to create the final construct.

(6) pLAS2w.STK40-KD(50-300aa) only-P2A-EYFP compries only the kinase domain of STK40, which is the amino acids 50 to 300 of STK40. pLAS2w.STK40-P2A-EYFP was digested with restriction enzymes NheI and SbfI with the insert STK40-KD ligated to it. STK40-KD was generated by performing in-fusion PCR cloning utilizing primers *STK40 35 ATG infusion F* and *STK40 331 infusion R*, Phusion DNA polymerase (Thermo Fisher Scientific) and the template pLAS2w.STK40-EYFP.

To create a cell line SAS-GFP-PXN for FA dynamics, we created pLKO.AS3W-GFP-PXN.bsd.

(7) pLKO.AS3W-GFP-PXN.bsd was created by digesting vector pLKO.AS3w.bsd (Academia Sinica) using restriction enzymes NheI and SbfI, then ligating cloned GFP-PXN to the vector. GFP-PXN was cloned utilizing primers *GFP-PXN-F* and *GFP-PXN-R* and template GFP-PXN (gift from Tobias Meyer's lab). pLKO.AS3W-GFP-PXN.bsd was later expressed in SAS cells and sorted to create cell line SAS-GFP-PXN to label focal adhesions in FA dynamic experiments.

Lentivirus preparation and infection

HEK293T cells were plated on 6 cm or 10 cm dish (JET Biofil ®, Guanzhou, China) and transfected with shRNA plasmid or cloned construct, pMD2.G and pCMV Δ8.91 (gifts from Ching-Chow Chen's lab, National Taiwan University) using Lipofectamine ® 3000 (Invitrogen, Thermo Fisher Scientific, Waltham, MA, USA) and P3000 enhancer (Invitrogen) in OPTI-MEM (Gibco). After 6 hr of transfection, the medium was removed and replaced with DMEM with 1% BSA. Viral supernatant was collected for 48 hr after transfection, centrifuged and concentrated using Lenti-X (ClonTech, Takara, Shiga, Kyoto, Japan). SAS, HUVEC, or HepG2 cells were infected with lentiviruses, including 8 μg/mL polybrene (SantaCruz, Santa Cruz, CA, USA) for enhancement of infection. Cells were then selected using 2 μg/mL puromycin (Sigma, St. Louis, MI, USA) for shRNAs or 10 μg/mL blasticidin (InvivoGen) for overexpression constructs for 24 to 48 hr.

"Two-hit" migration screen

High Throughput Screening (HTS, High content screening, HCS) service and shRNA provided by National RNAiCore (NRC) in Academia Sinica was utilized to accelerate the process of the "two-hit" migration screen. HUVEC cells were plated manually on 96 well plates (Corning, Costar 3599) coated with 30 μg/mL collagen I (Rat Tail, Invitrogen). 119 genes were selected from two previous screens (Vitorino et al., 2008; Simpsons et al., 2008) as our candidates in the "two-hit" migration screen. A total of 588 shRNAs targeting the 119 genes were used to infect HUVEC cells operated by robotic arms. On the day of wound healing assay, HUVEC cells were stained with Hoechst 33342 (Invitrogen) in EGM2 medium at room temperature. After nuclei staining, the medium was changed to EGM2 containing DMSO (Sigma), 2 μM BTP2 (Sigma), 5 μM Y27632 (Sigma), or 10 μM PD98059 (Sigma). A scratcher (Vitorino et al., 2008) was then used to create wounds for wound healing migration. Plates were sent to ArrayScar ® VTI HCS Reader (Thermo Fisher Scientific, Cellomics$^{TM}$) to take images at 0 hr and 15 hr of incubation at 37°C.

Data from our "two-hit" migration screen were analyzed utilizing Matlab (MathWorks, Natick, MA, USA) in the following flow (Data S5: *HUVEC sheet migration analysis script*):
(1) Generation of wound healing area (2) Density correction (3) Normalization (4) Z-score

(1) The wound-healing area of the sheet migration images between 0 hr and 15 hr in the "two-hit" migration screen on HUVEC cells were analyzed. (Data S1, *sheet: Wound healing area*). The extent of wound healing after wound-scratch was demonstrated as the wound healing area.

(2) Density correction (fig. S1B) was carried out by generating a linear regression curve of sheet migration versus cell density. The estimated sheet migration per density was generated based on the regression line. Density corrected speed was calculated as the following formula:

$$Density\ corrected\ speed = actual\ speed - estimated\ speed$$

(3) Normalized sheet migration was calculated as the following formula. (Data S1, *sheet: Normalized wound healing*, Fig. 1D).

$$Normalized\ sheet\ migration\ = \frac{Sheet\ migration\ of\ each\ shRNA}{average\ sheet\ migration\ of\ untreated\ cells}$$

(4) Z-scores of sheet migration was calculated as the following formula (Data S1, *sheet: Z-score*, fig. S1, C and D):

$$z - score = \frac{(wound\ healing\ area\ of\ the\ gene - average\ wound\ healing\ area\ of\ all\ shRNAs\ under\ treated\ vehicle)}{standard\ deviation\ of\ the\ wound\ healing\ area\ of\ all\ shRNAs\ under\ treated\ vehicle}$$

Positive z-score values imply greater wound healing while negative values imply less healing. Take STK40 as an example: the z-score of the N column of shSTK40 is close to 0, while shSTK40 plus PD98059 generated a negative value. This implies a synergistic reduction in sheet migration when co-inhibiting STK40 and MAPK.

Single-Cell Tracing Migration (Random migration and Wound healing assays)

SAS and HUVEC cells were plated on collagen I (Rat Tail, Invitrogen) coated 96 well plates (Corning, CoStar 3599). Cells were infected with lentiviruses for 24 hr and selected with antibiotics for 24 to 48 hr (see *Lentivirus preparation and infection*). On the day performing migration assays, SAS cells were stained with 1 µg/mL of Hoechst 33342 for 1 hr at 37°C, while HUVEC cells were stained with 1 µg/mL of Hoechst 33342 for 15 to 30 min at the same temperature. After nucleus staining, medium of SAS cells was changed to serum-free medium containing 20 mM HEPES (Gibco), 0.1% BSA (BioShop, Ontario, Canada), and 0.5 ng/mL EGF (PeproTech, Rockyhill, NJ, USA), or supplemented with 25 ng/mL FGF1 (Invitrogen) plus 10 U heparin (Sigma). Medium of HUVEC cells were changed to EGM2 medium. Addition of other drugs such as trametinib 100 nM (LC laboratories, Woburn, MA, USA) or DMSO was added into the medium together with the supplements. For wound healing assays, cells were scratched by a scratcher while random migration assays do not require wound scratch. After wound-scratch (or un-scratched), the cells were then placed in an acrylic incubator at 32°C for SAS cells and 37°C for HUVEC cells, utilizing Nikon Eclipse *Ti* microscope (Nikon, Tokyo, Japan) for live-cell imaging. Images were taken every 10 min for a duration of 6 to 10 hr. Parameters such as speed, coordination and directionality were analyzed using Matlab (Data S5: *Cell migration assay scripts*).

Double-inhibition Experiment

SAS cells were plated on collagen I (Rat Tail) coated 96 well plates. For double-inhibition experiments, cells were co-inhibited with (1) shSTK40 plus (2) shRNAs or drug targeting migration related molecules: Rac1 for F-actin, α-catenin, for adherens junction, paxillin, focal adhesion, and myosin light-chain kinase for myosin. Cells were infected with shSTK40 plus shCTNNA1, shRAC1, or shPAXILLIN (shPXN) for 24 hr and selected with puromycin for 24 hr before performing migration. 10 µM of ML9 (Sigma), myosin-light chain kinase inhibitor, was added into serum free DMEM medium with supplements after nucleus staining on the day performing migration. Random migration or wound-healing migration were carried out after infection and selection (see *Single-Cell Tracing Migration*). Nucleus stained or drug treated SAS cells, scratched or unscratched were then placed in an incubator at 32°C for migration, utilizing Nikon Eclipse *Ti* microscope for live-cell imaging. Images were taken every 10 min for a duration of 10 hr followed by analysis using Matlab.

Immunofluorescent staining, IF

SAS , HUVEC, and HepG2 cells were plated on 100 µg/mL collagen (Collagen I, bovine, Gibco) coated chamber slides (Thermo Fisher Scientific, 155411 or 155383 Nunc™ Lab-TEK™) or 96 well plate (Thermo Fisher Scientific, Nunc™ 165305) and infected with shRNA viruses or prepared viruses of overexpression constructs followed by aforementioned antibiotic selection.

SAS cells were treated with or without drugs before fixation. Drugs include 5 µM of Y27632, 5 µM of blebbistatin, 100 nM of trametinib treatment overnight, 2.5 µM verteporfin (MedChemExpress, Monmouth Junction, NJ, USA) 8 hr, and 100 nM leptomycin B (Cayman Chemical Company, Ann Arbor, MI, USA) 2 to 3 hr.

Prior to fixation, cells were processed to a 2 hr-migration. Cells were washed with PBS and wounds were created using a tip for chamber slides or a scratcher for 96 well plates. The cells were incubated in serum-free DMEM supplemented with 20 mM HEPES, 0.1% BSA, 25 ng/mL FGF1 and 10 U heparin, including or excluding drug treatment at 37°C for 2 hr to perform migration. After 2 hr of migration, cells were fixed with 4% paraformaldehyde (Sigma) in PBS at room temperature for 15 min, permeated with 0.25% Triton-X-100 (J.T.Baker, Phillipsburg, NJ, USA) at room temperature for 10 min and blocked with 5% BSA at room temperature for 1 hr. Cells were then incubated overnight at 4°C with primary antibodies with 1% BSA in PBS. The cells were then stained with secondary antibodies (see *Antibodies*), phalloidin dyes, and 10 µg/mL of DAPI, (Invitrogen) with 1% BSA in PBS. Phalloidin dyes, Alexa-Fluor 488 ® phalloidin (Invitrogen, Cat #A12379, 1:500 to 1:1000 dilution) and Alexa-Fluor 594 ® phalloidin (Invitrogen, #A12381,1:500 to 1:1000 dilution), were used to stain stress fibers. We then utilized Nikon Eclipse *Ti* to take images of the fixed cells.

Western blotting

Cells were infected with viruses and treated with or without drugs such as PD98059, trametinib, Y27632, and blebbistatin (Sigma) for 24 hr. Cells were washed in ice-cold PBS and then harvested in radioimmunoprecipitation assay (RIPA) lysis buffer (Cell Signaling Technology, Danvers, MA, USA) and supplemented with phenylmethylsulfonyl fluoride (PMSF) (MD BioInc., Rockville, MD, USA) and protease and phosphatase inhibitor cocktail (Thermo Fisher Scientific). Lysates were centrifuged for 15 min at 15000 rpm. Samples that were boiled at 95°C for 5 min were loaded into wells of sodium dodecyl sulfate polyacrylamide gel electrophoresis (SDS-PAGE) gel, transferred onto polyvinylidene difluoride (PVDF) membrane (Merck Millipore, Burlington, MA, USA, Immobilon ®-P PVDF Membrane), blocked with 3% BSA in 0.05% TBST (BioLink, Lisle, IL, USA) and incubated at 4°C overnight with primary antibodies and secondary antibodies at room temperature for 1 hr (see *Antibodies*). Enhanced chemiluminescent (ECL) substrates (T-Pro Biotechnology, New Taipei City, Taiwan) was used to visualize conjugated proteins, followed by imaging using Bio-Rad Gel Doc2000 (Bio-Rad, Hercules, CA, USA). Image Lab Software (Bio-Rad) was utilized for protein quantification.

<u>Antibodies</u>

Antibodies in IF were as follows: Purified mouse anti-paxillin (BD BioSciences, San Jose, CA, USA, Cat #610061 and #610062) at 1:150; Phospho-Myosin Light Chain 2 (Ser19) Antibody (Cell Signaling Technology, Cat #3671) at 1:1000; YAP (D8H1X) XP® Rabbit mAb (Cell Signaling Technology, Cat #14074) at 1:1000; Goat anti-Mouse IgG (H+L) Cross-Adsorbed Secondary Antibody, Alexa Fluor 488 (Invitrogen, Cat #A11001) at 1:500; Goat anti-Mouse IgG (H+L) Cross-Adsorbed Secondary Antibody, Alexa Fluor 594 (Invitrogen, Cat #A11005) at 1:500; Goat anti-Rabbit IgG (H+L) Cross-Adsorbed Secondary Antibody, Alexa Fluor 488 (Invitrogen, Cat #A11008) at 1:500, Goat anti-Rabbit IgG (H+L) Cross-Adsorbed Secondary Antibody, Alexa Fluor 594 (Invitrogen, Cat #A11012) at 1:500.

Antibodies in Western blots were as follows: Phospho-p44/42 MAPK (Erk1/2) (Thr202/Tyr204) Antibody (Cell Signaling Technology, Cat #9101) at 1:5000; p44/42 MAPK (Erk1/2) Antibody (Cell Signaling Technology, Cat #9102) at 1:5000; Phospho-MEK1/2 (Ser217/221) (41G9) Rabbit mAb (Cell Signaling Technology, Cat #9154) at 1:1000; MEK1/2 (L38C12) Mouse mAb (Cell Signaling Technology, Cat #4694) at 1:1000; Phospho-c-Raf (Ser259) Antibody (Cell

Signaling Technology, Cat #9421) at 1:1000; c-Raf Antibody (Cell Signaling Technology, Cat #9422) at 1:1000; Phospho-FAK (Tyr397) Antibody (Cell Signaling Technology, Cat #3283) at 1:1000; FAK Antibody (Cell Signaling Technology, Cat #3285) at 1:1000; Phospho-YAP (Ser127) (D9W2I) Rabbit mAb (Cell Signaling Technology, Cat #13008) at 1:1000; YAP (D8H1X) XP® Rabbit mAb (Cell Signaling Technology, Cat #14074) at 1:1000; α-Tubulin (DM1A) Mouse mAb (Cell Signaling Technology, Cat #3873) at 1:5000; E-Cadherin antibody [N3C2], Internal (GeneTex, Irvine, CA, USA, Cat #GTX124178) at 1:1000; Anti-alpha smooth muscle Actin antibody (Abcam, Cambridge, UK, Cat #ab5694) at 1:1000 dilution; GAPDH Loading Control Monoclonal Antibody (GA1R) (Thermo Fisher Scientific, Cat #MA5-15738) at 1:5000. HRP conjugated Goat anti-Rabbit IgG (H+L) Mouse serum adsorbed (Abgent, San Diego, CA, USA, Cat #LP1001c) at 1:2000; HRP conjugated Goat anti-Mouse IgG (H+L) Human serum adsorbed (Abgent, Cat #LP1002c) at 1:2000.

Real-time reverse transcription polymerase chain reaction (RT-qPCR)

Total RNA of cells were extracted using Iso-RNA lysis reagent (Five Prime Therapeutics, South San Francisco, CA, USA) and converted into cDNA using SuperScript Transcriptase II (Invitrogen). Samples reactions containing SYBR Green Supermix (Bio-Rad), primers (Data S4) and cDNA template were then loaded onto 96 well PCR plates (Bio-Rad) then set up for real-time PCR using CFX Connect Real-Time PCR Detection System (Bio-Rad). Matlab was utilized to analyze PCR cycles, annealing temperature and quantification of mRNA levels (Data S5: *QPCR*).

Focal adhesion analysis

Immunofluorescent images of focal adhesion were analyzed using Matlab (Data S5: *FA scripts*). FA quantification was performed using the following flow: (1) FA background subtraction: Script *FA_background_20200313.m* was utilized for FA background subtraction. The background signal of each of the individual pixels was separately determined by the median value of its surrounding pixels within the radius of 1.46 μm. (2) Thresholding: Script *FA_identification_FAclearer_20200316.m* was utilized. The threshold for the mask of FA was determined as two standard deviations above the mode of the background value. (3) Segmentation: Function of watershed segmentation in Matlab was used for FA segmentation. (4)

FA selection: FAs were identified and selected in a semi-automatic manner. FAs within 15 μm of the lamellipodia edge were further analyzed. For each FA, we acquire its area and mean FA signal. The integrated FA signal (fig. S4E) was generated by:

$$Integrated\ FA = FA\ area \times mean\ FA\ signal$$

FAs and cells from multiple sites were analyzed and data were pulled up using Microsoft Excel (Redmond, WA, USA) for statistical analysis. Here is a list of the cells and FAs chosen in each figure:

1. FAs of 8 cells at the wound edge from 1 site of each condition were analyzed for STK40 knockdown and overexpression (Fig. 2, D and E) in SAS cells. Number of FA per cell was also analyzed (fig. S5A). FAs of 23 cells at wound edge from 1 site in cells infected with shControl and 31 cells from 1 site in cells infected with shSTK40 were analyzed in HepG2 cells (fig. S5B). FAs of 14 cells at the wound edge from 1 site in cells infected with shControl and 13 cells from 1 site in cells infected with shSTK40 were analyzed in HUVEC cells (fig. S5C).
2. FAs of 8 cells at the wound edge from 1-2 sites of each treatment were analyzed for the overexpression and kinase domain truncation of STK40 (Fig. 3, A and B).
3. FAs of 24 cells at the wound edge from 3 sites of each condition were analyzed in the experiment translocating STK40 from nucleus to cytosol (Fig. 3, C and D).
4. FAs of 8 cells at the wound edge from 1 site of each treatment were analyzed in the experiment removing traction force upon knockdown of STK40 (Fig. 3, F and G).
5. FAs of 8 cells at the wound edge from 1 site of each condition were analyzed in the experiment of shSTK40 plus trametinib (Fig. 4A).
6. FAs of 8 cells at the wound edge from 1 site of each condition were analyzed in the experiment of shSTK40 plus verteporfin (Fig. 4C).
7. FAs of 8 cells at the wound edge from 2-4 sites of each condition were analyzed in the experiment of shSTK40 plus trametinib and verteporfin (Fig. 4D).
8. FAs of 8 cells at the wound edge from 1 site for each condition were analyzed in the experiment of shSTK40 plus leptomycin (Fig. 4F).

9. FAs of 8 cells at the wound edge from 1 site for each condition were analyzed in the experiment of shSTK40 plus verteporfin and shCREB (Fig. 4G).

Images of FA shown in figures were processed utilizing Matlab (Data S5: *FA demo*). Script *Generate_FA_demo_20200831.m* was utilized.

Focal adhesion dynamic assay

Sorted cells were used to create cell line SAS-GFP-PXN for FA dynamics experiments. We infected SAS cells with viruses packaged pLKO.AS3w-GFP-PXN.bsd and cultured the cells for cell sorting. After trypsinizing and centrifuging infected cells, the supernatant was removed and cells were treated and resuspended with 1mL of sorting buffer that includes 1X PBS ($Ca^{2+}$ / $Mg^{2+}$ free), 1 mM EGTA (Sigma), 2 mM HEPES, 1-2% FBS and 0.1 to 0.2% BSA, at a pH of 7.0. Cells were filtered and prepared with sterilized sorting tube (BD Biosciences, BD falcon 352235) before cell sorting using flow cytometer BD LSRFortessa™ (The Flow Cytometric Analyzing and Sorting Core of the First Core Laboratory, College of Medicine, National Taiwan University).

96 well with coverglass base plates (Thermo Fisher Scientific, Nunc™ 164588) were coated with 100 µM poly-D-lysine hydrobromide (Sigma) for 5 min of incubation at 37°C and then coated with 10 µg/mL collagen (Collagen I, Rat tail) overnight. GFP-PXN-SAS cells were then plated for FA dynamic assay. On the day of FA dynamic assay, the medium of cells was removed and replaced with phenol red-free serum free medium DMEM (Gibco) supplemented with 10% of fetal bovine serum and 100 µM L-ascorbic acid (Sigma, Cat #A4544). Movies of live-cells were recorded using Nikon Eclipse *Ti* microscope with a frequency of 5 sec per image and a total duration of 8 min. FA dynamics were analyzed using Matlab (Data S5: *FA dynamics*). All FAs of all cells in 1 site of each treatment (shControl or shSTK40) were analyzed.

Flow cytometry

SAS cells were knocked down with shControl and shSTK40. Samples were trypsinized and centrifuged, followed by resuspension with a staining buffer: 1% fetal bovine serum, (Biological Industries, Beit-Haemek, Israel), 2 mM of EDTA (J.T. Baker) and 0.05% sodium azide (Sigma) in PBS on ice. Flow cytometry was performed on flow cytometer BD LSRFortessa.

YAP analysis

Immunofluorescent images of YAP were analyzed using Matlab (Data S5: *YAP*). Script *N_C_calculation_20190627_YFP_20191026_YAP.m* was utilized for YAP analysis. To determine YAP signals of a single cell, the image of its DAPI signal was used as a nuclear mask. Its nuclear YAP signals were determined by the YAP signals overlying the nuclear mask. Its cytosolic YAP signals were determined by the YAP signals on the ring area of 0.73 μm in width surrounding the nuclear mask. YAP levels of shControl or shSTK40 treated cells were analyzed from 12 cells in 1 site (Fig. 4E).

Statistical Analysis

Experimental data such as bar graphs, error bars are presented as mean ± SE (standard error). Two-tailed Student's *t* tests were performed with Microsoft Excel and Matlab. Differences were considered statistically significant when *P* values were less than 0.05.

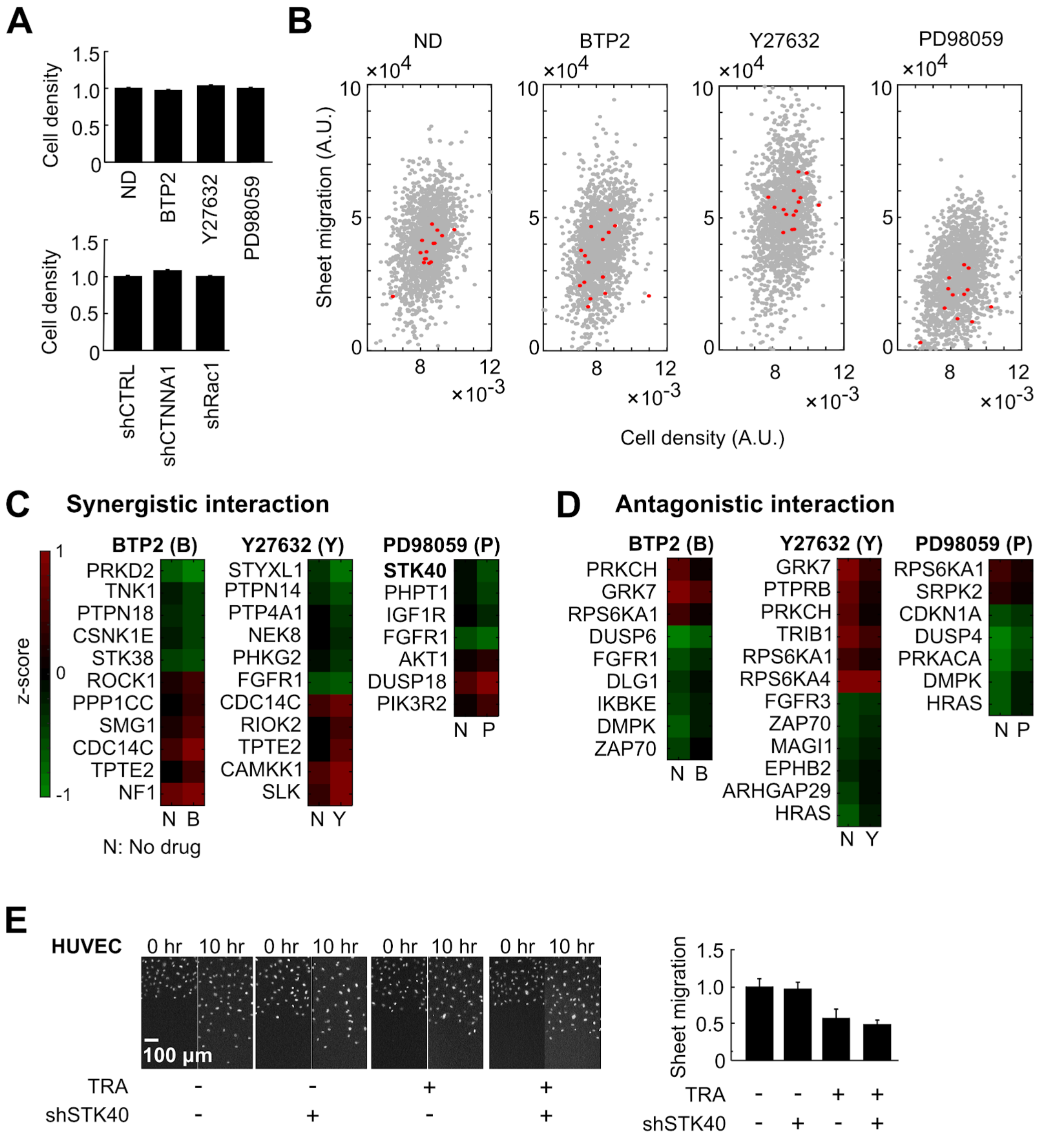


**Fig.S1**

**Fig. S1. A two-hit screen revealed signaling interactions during sheet migration.**

(A to D) Data of the two-hit screen in HUVECs. (A) Density of cell sheets under small molecule inhibitors (upper panel) or shRNAs (lower panel). Concentrations of drugs (BTP2, Y27632 and PD98059) and viral titers of shRNAs (shCTRL, shCTNNA1 and shRAC1) were titrated to avoid significant changes of cell densities. ND: no drug. (B) Scatter plots demonstrate effects of shRNAs on sheet migration versus cell density under different inhibitors. Red dots mark shSTK40s. Density correction was conducted based on linear regression using these dots. (C and D) Analysis of synthetic dysmobility. Normalized and density-corrected sheet migration values of individual shRNAs converted to z-scores using data of all shRNAs under the same inhibitor. (C) For each shRNA, the interaction was considered synergistic if the magnitude of its z-score was increased by the inhibitor, or (D) antagonistic if the magnitude was decreased. (E) The shSTK40-MAPK inhibition was verified using another MAPK inhibitor trametinib (TRA). Left:

Representative images. Right: Quantification. Error bars denote mean ± standard error of the mean (s.e.) from six biological replicates.

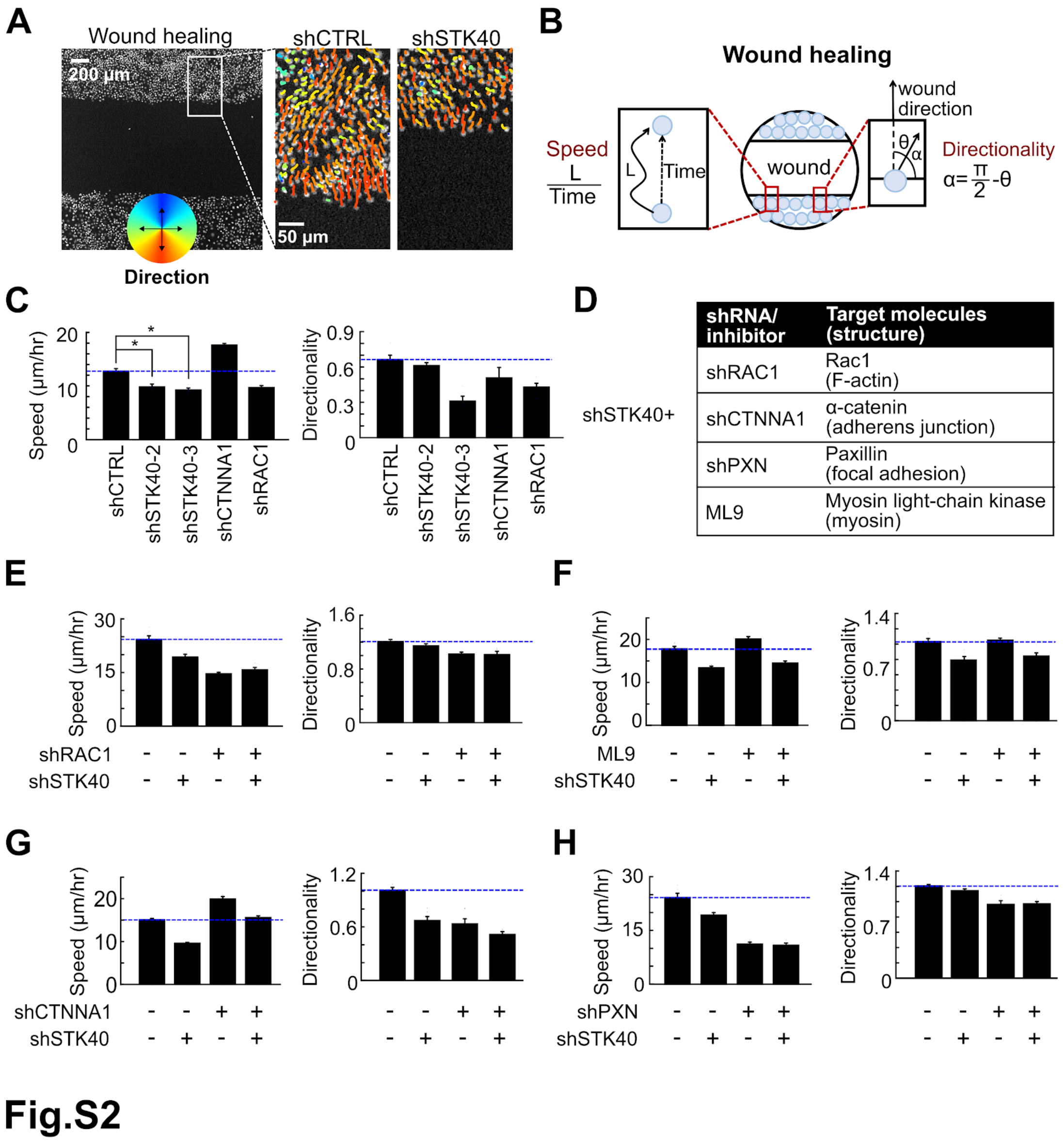


**Fig. S2. Exploration of STK40 effects on wound-healing migration using single-cell tracing.** (A) (Left) Cell nuclei were labeled with Hoechst 33342. (Right) Colored lines depicted traces and directions of migration from individual cells treated with shCTRL or shSTK40 within the two-hour period. (B) Cell speed and directionality were used to evaluate wound healing migration. (C) Compared to random migration (Fig. 2B), shSTK40 had less suppressive effect on cell speed and inconsistent effect on directionality. **P* < 0.05. (D) Design of double-inhibition experiments. (E to H) Results of double-inhibition experiments in wound healing assays. SAS cells were infected with shSTK40 and / or (E) shRAC1, (F) ML9, (G) shCTNNA1 and (H) shPXN. Error bars denote mean ± s.e. from six biological replicates.

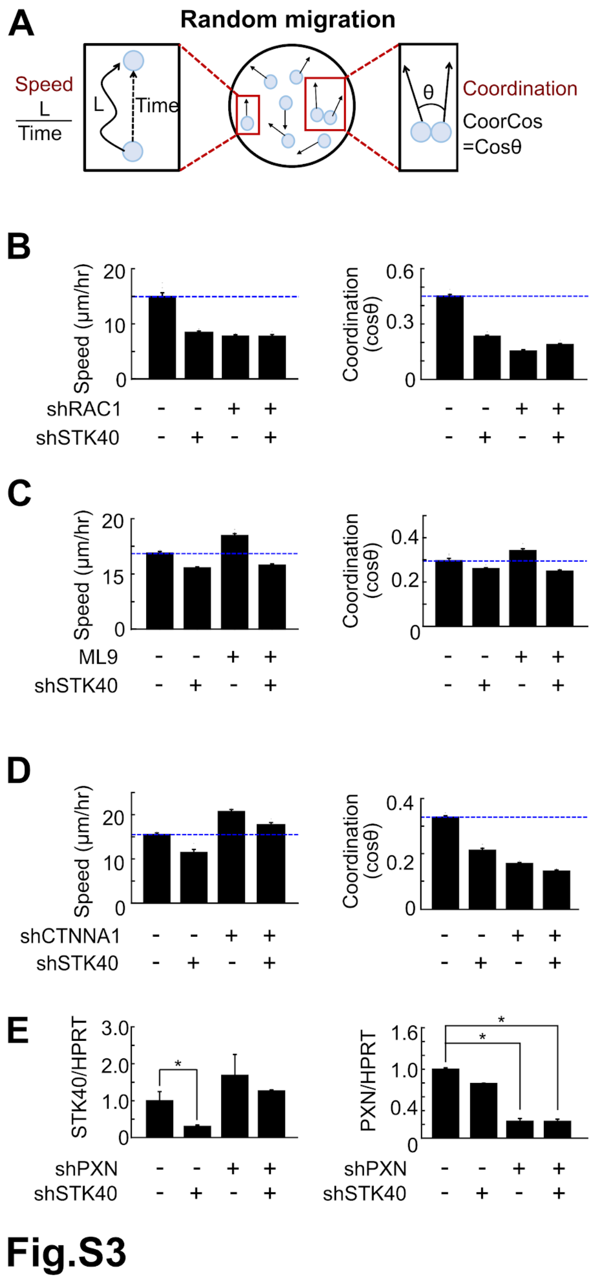


**Fig. S3. Exploration of STK40 effects in random migration using double-inhibition experiments.**

(A) Cell speed and coordination were used to evaluate random migration. (B to D) SAS cells were co-inhibited with shSTK40 and / or (B) shRAC1 (C) ML9 and (D) shCTNNA1. (E) Knockdown efficiency of shSTK40 and shPXN was evaluated by measuring mRNA levels of STK40 and paxillin (PXN) using RT-qPCR. Error bars denote mean ± s.e. from six biological replicates in (B to D) and three biological replicates in (E). *$P < 0.05$.

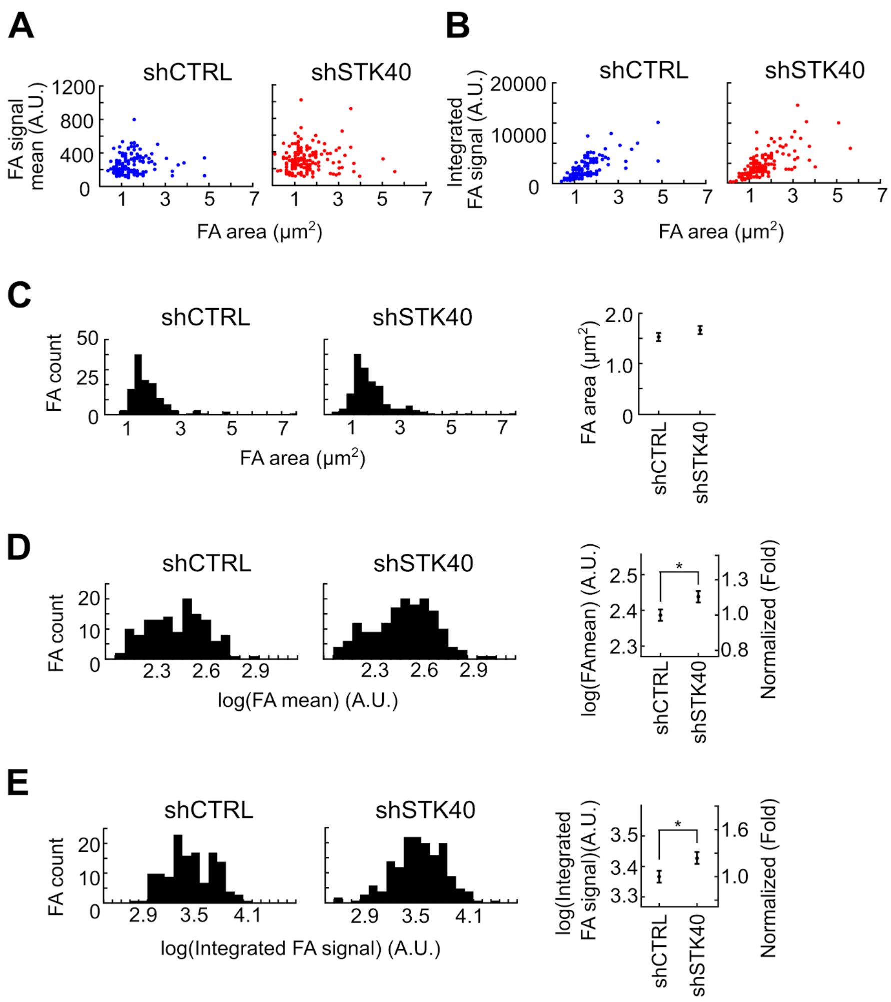


**Fig.S4**

**Fig. S4. STK40 knockdown increased the magnitude of focal adhesion complexes.** Immunofluorescence using anti-paxillin antibody was used to label focal adhesion (FA) complexes. Area and mean FA signals were measured for individual FAs. (A) Scatter plots of area versus mean FA signals for individual FAs. (B) Scatter plots of area versus integrated FA signals, generated by (FA area)*(mean FA signal). (C) Histograms (left) and error bar (right) graph reveal a small increase of FA area by shSTK40 compared to shCTRL. (D) Histograms (left) and error bar (right) graph reveal a remarkable increase of the logarithm of mean FA signals by shSTK40 compared to shCTRL. (E) Histograms (left) and error bar (right) graph reveal a remarkable increase of the logarithm of integrated FA signals by shSTK40 compared to shCTRL. Note that normalized values from the logarithm of integrated FA signals were used to represent FA magnitudes as “FA (fold)” in all following figures (Fig. 2, Fig. 3, Fig. 4, fig. S5, fig. S6, and fig. S7). Error bars denote mean ± s.e. $*P < 0.05$.

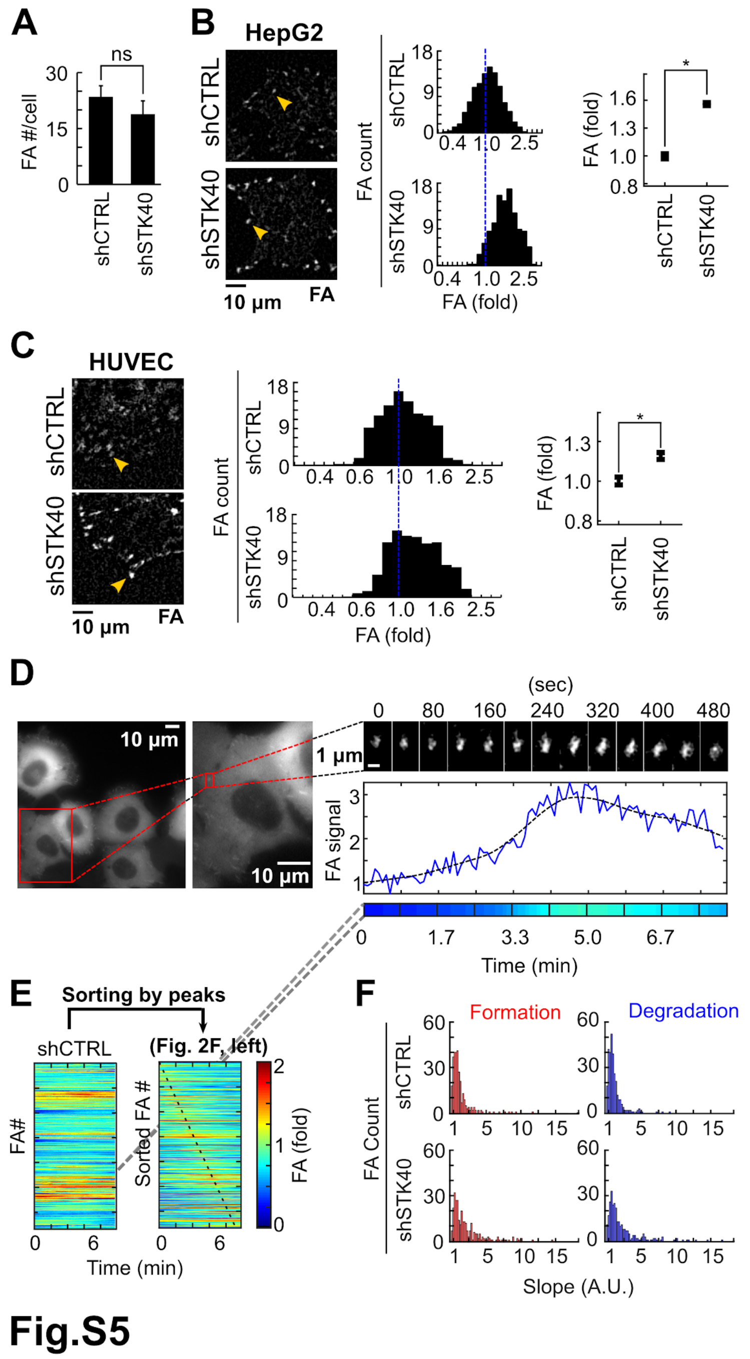


**Fig. S5. Knockdown of STK40 increased focal adhesion in different cell lines by altering focal adhesion dynamics.**

(A) Quantification of numbers of focal adhesion (FA) complexes per cell. shSTK40 did not increase the number of FAs. (B and C) (Left) Images of FA upon shSTK40 treatment in (B) HepG2 and (C) HUVEC cells. Their integrated FA signals were quantified as histograms (middle) and error bar graph (right). Error bars denote mean ± s.e. **P* < 0.05. (D to F, together with Figure 2, F to I) Analysis of FA dynamics based on time lapse images. (D) Left: SAS cells expressing GFP-PXN were used for the analysis. Right: Kymograph of signals of chosen FA over time. Its quantification was plotted as the blue line and smoothened as the dotted line. Color bar depicts the signals using the color scale in (E). (E) Color bars as rows from individual FAs were stacked to generate the color map (left). Then the rows were sorted based on the timing of their peaks to generate the color map in Figure 2F. (F) For each FA, rates of its formation and degradation were calculated as derivatives of signals by time (presented as “Slope”) before and after the signal peak. Histograms depict slopes of FA formation and degradation from individual FAs.

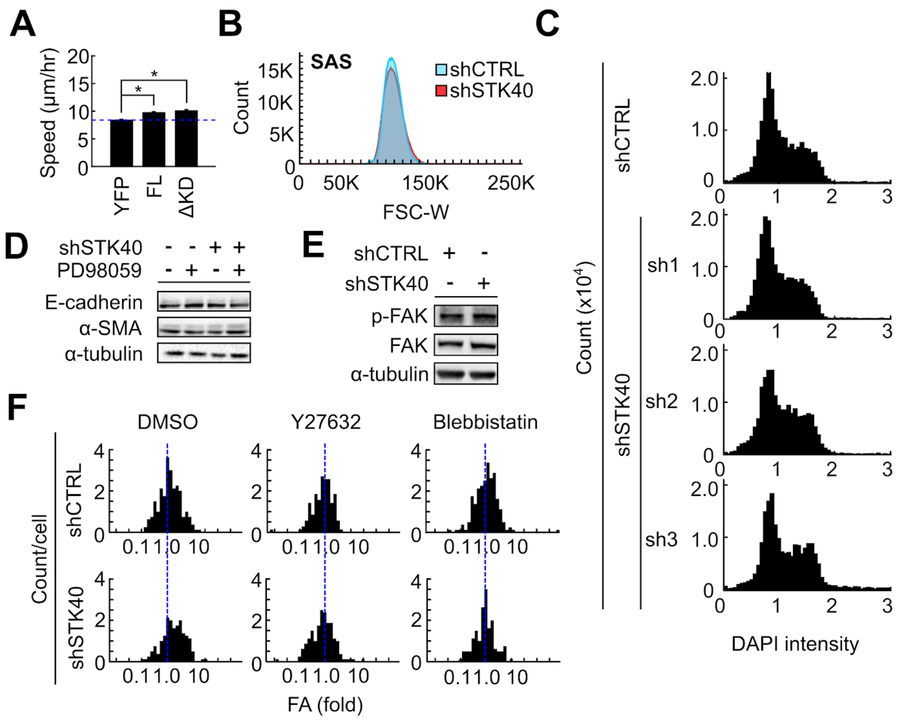


**Fig.S6**

**Fig. S6. STK40 did not alter focal adhesion through its kinase activity, cell proliferation or differentiation, nor focal adhesion kinase signaling.**

(A) Speed of SAS cells over-expressing empty vector (YFP), STK40 of full-length (FL), and STK40 with kinase-domain truncated (ΔKD) in random migration. Notice that ΔKD rescued cell migration similarly to FL. $*P < 0.05$. (B) Flow cytometry showed that shSTK40 did not alter cell size. (C) Using DAPI to label DNA contents of individual cells and depicting their DAPI intensities in histograms, we demonstrated that shSTK40 did not alter DNA contents, indicating its minor effect on cell cycles. (D) Western blots showed that shSTK40 or MAPK inhibitor PD98059 had minimal effects on E-cadherin or α-SMA levels, indicating the minor role of STK40 on the epithelial-mesenchymal transition. (E) Western blots showed that shSTK40 did not change the phosphorylation status (p-FAK) of focal adhesion kinase (FAK). (F) Histograms of FAs in SAS cells infected with shSTK40 and treated with drugs (DMSO, Y27632 5 μM, or blebbistatin 5 μM). Notice that the rightward shift of FA distribution by shSTK40 was abolished by Y27632 or blebbistatin, indicating the importance of myosin-mediated traction force on STK40-mediated FA strengthening. The results were further quantified as error bars in Figure 3G.

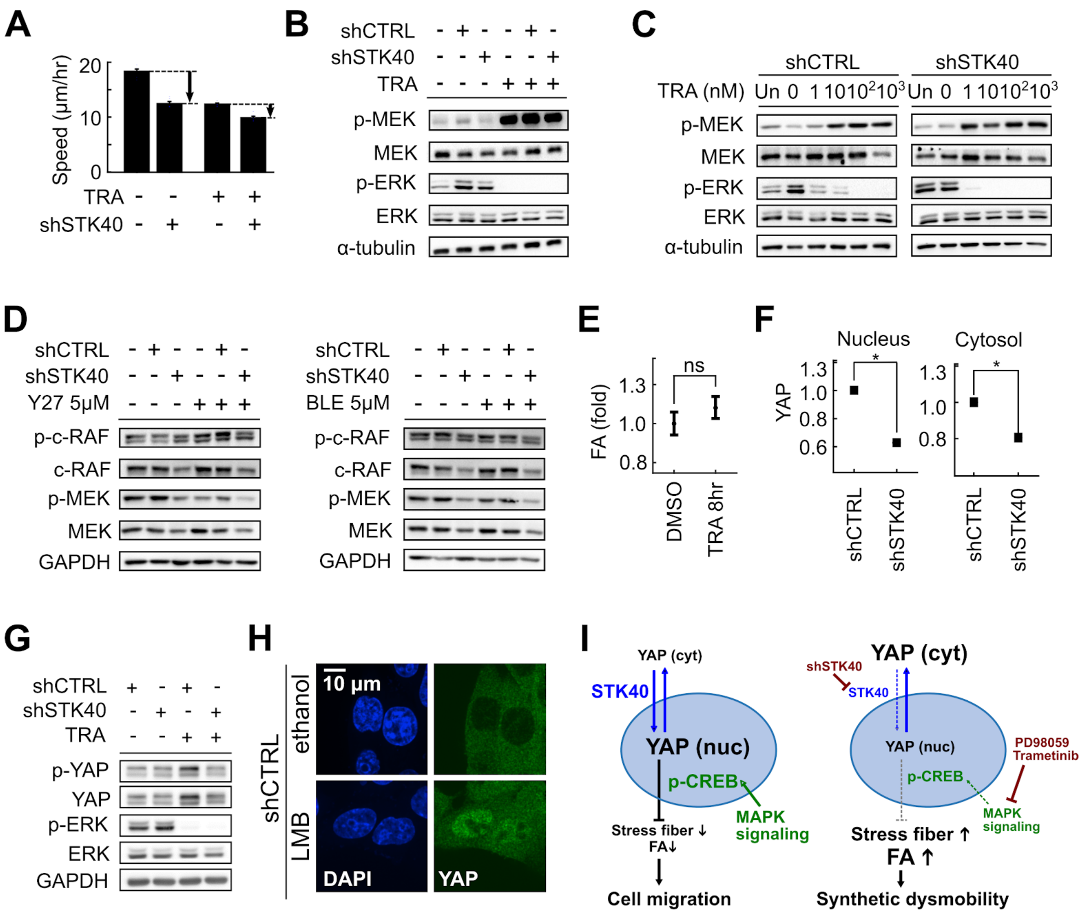

Fig.S7

**Fig. S7. STK40 collaborated with MAPK signaling to regulate YAP-mediated FA remodeling.**

(A) Speed of SAS cells with shSTK40 and / or MAPK inhibitor trametinib (TRA) in wound healing migration. (B) Western blots revealed that shSTK40 moderately reduced phosphorylation of MEK and ERK. In contrast, trametinib completely suppressed ERK phosphorylation with compensatory enhancement of MEK phosphorylation. (C) Western blots showed that trametinib concentration-dependently suppressed ERK phosphorylation with compensatory MEK phosphorylation, both of which were sensitized by shSTK40. (D) Western blots showed that shSTK40 did not reduce phosphorylation of c-RAF but mildly decreased its protein level. ROCK inhibitor Y27632 (Y27) or myosin inhibitor blebbistatin (BLE) did not alter shSTK40 effects on MAPK signaling. (E) Error bars depict that trametinib treatment for 8 hours had not significantly changed FA. ns, not significant. (F) Error bars show that shSTK40 reduced nuclear and cytosolic YAP levels. (G) Western blots showed that shSTK40 slightly reduced YAP levels but completely reversed the accumulation of YAP due to compensatory responses under MAPK inhibition. (H) Immunofluorescence shows that leptomycin-B (LMB) caused YAP accumulation in cell nuclei. (I) Working models depict how STK40 and MAPK collaboratively regulate YAP activities (left) and how STK40 plus MAPK suppression causes “synthetic dysmobility.” (right). Error bars denote mean ± s.e. *$P < 0.05$.

**Movie S1. Random migration of SAS cells under shCTRL.**
Movie of random migration in SAS cells with shCTRL (Fig. 2A). Cell nuclei were labeled with Hoechst 33342. Traces of cell migration were added by computation using Matlab. Color pie chart reveals directions of cell traces.

**Movie S2. Random migration of SAS cells under shSTK40.**
Movie of random migration in SAS cells with shSTK40 (Fig. 2A). Cell nuclei were labeled with Hoechst 33342. Traces of cell migration were added by computation using Matlab. Color pie chart reveals directions of cell traces.

**Movie S3. Wound-healing assay in SAS cells under shCTRL.**
Movie of wound-healing assay in SAS cells with shCTRL (fig. S2A). Cell nuclei were labeled with Hoechst 33342. Traces of cell migration were added by computation using Matlab. Color pie chart reveals directions of cell traces.

**Movie S4. Wound-healing assay in SAS cells under shSTK40.**
Movie of wound-healing assay in SAS cells with shSTK40 (fig. S2A). Cell nuclei were labeled with Hoechst 33342. Traces of cell migration were added by computation using Matlab. Color pie chart reveals directions of cell traces.

**Movie S5. Focal adhesion dynamics.**
Demonstration of the focal adhesion (FA) dynamics and quantitative analysis (fig. S5D). SAS cells expressing GFP-PXN were used for live-cell fluorescence imaging. Red boxes denote the specific cell and FA chosen for the analysis of FA dynamics. Quantification of FA signals over time was plotted as the blue line. Dotted line represents smoothened signals.

**Data S1. Original data of "two-hit" migration screen: wound healing area, normalized wound healing area, and z-scores.**
Sheet 1 (Original): Original wound healing area (A.U.) of all shRNAs used in the "two-hit" migration screen. Name of the genes, Clone ID of shRNAs, vector of shRNA, region that shRNA knocked down, and complete names of genes were listed. Three replicates of untreated (yellow), BTP2 (red), Y27632 (blue), and PD98059 (green) treated were shown.

Sheet 2 (Normalized): Data of original wound healing area were normalized to the average wound healing area of all shRNAs under treated vehicles (DMSO, BTP2, Y27632 or PD98059).

Sheet 3 (Z-scores): The z-scores of all shRNAs in the "two-hit" screen generated by the formula mentioned in *"Two-hit"migration screen* in *Materials and Methods*. Positive values resemble greater wound healing while negative values symbolize less healing.

**Data S2. List of shRNAs and targeting sequences utilized for experiment.**
Chart of the shRNAs utilized to create viruses to knock down particular genes. The cloneID,

targeting gene region (3'UTR or coding DNA sequence, CDS), and targeted gene sequences for knockdown were listed.

**Data S3. List of templates, inserts, primer sequences, and restriction enzymes utilized to create overexpression constructs.**

Chart of plasmids of the overexpression constructs utilized in paper. Templates, vectors, cloning inserts, sequence of primers, and restriction enzymes utilized to create constructs were listed.

**Data S4. qPCR primers targeting different genes utilized for Real-Time PCR experiment.**

Chart of the qPCR primers utilized in paper. Primer names and sequences are shown in the table.

**Data S5. Matlab codes utilized for image processing and analysis.**

All Matlab codes utilized for analysis for all experiments were listed.
